# Autophagy inhibition in intestinal stem cells favors enteroendocrine cell differentiation through Stat92E activity

**DOI:** 10.1101/2024.07.05.602279

**Authors:** Camille Lacarrière-Keïta, Sonya Nassari, Steve Jean

## Abstract

Because the intestinal epithelium faces many stresses, dysregulation of essential mechanisms governing gut homeostasis, such as autophagy, has been associated with inflammatory bowel pathologies. In *Drosophila melanogaster*, the inhibition of autophagy, specifically in adult intestinal stem cells (ISCs), affects their number differently through aging. Appropriate intestinal renewal requires a balance between ISC proliferation and differentiation. Herein, we show that in adult ISCs, the loss of core autophagy genes and regulators of autophagosome-lysosome fusion increased the enteroendocrine cell population and transcriptional activity of Stat92E. Functional experiments with cell fate regulators involved in enteroendocrine or enterocyte differentiation or proliferation suggested that dysfunctional autophagy in adult ISCs enhanced Stat92E activity downstream of Hop/JAK kinase. Finally, lineage-tracing analyses confirmed that autophagy inhibition autonomously promotes enteroendocrine cell differentiation without affecting enterocyte differentiation. Thus, our data demonstrates that, under homeostatic conditions, basal autophagy limits enteroendocrine cell differentiation by controlling Stat92E activity.

## INTRODUCTION

The intestinal tract faces multiple risks of tissue injury caused by food or pathogen ingestion and proximity to the microbiota^1^. Therefore, even under homeostatic conditions, rapid intestinal epithelium renewal is paramount to ensure adequate food digestion, nutrient absorption, and barrier function^2^. Intestinal cell turnover relies on intestinal stem cell (ISC) capacity to replenish damaged tissues with an adequate proportion of nutrient-absorbing and secretory cells^3^. External cues and inherent properties tightly regulate the fate of ISCs to balance proliferation and differentiation and preserve tissue functions^3^. Dysregulation of ISCs regenerative capacity can favor the development of inflammatory bowel disease (IBD)^4^.

Genome-wide association studies of IBD susceptibility genes have identified numerous autophagy regulators, suggesting that autophagy plays an essential role in intestinal homeostasis^5–7^. Accordingly, autophagy impairment in the intestinal epithelial cells leads to a loss of barrier integrity, failure to eliminate intracellular pathogens, decrease in antimicrobial defense, and microbiota dysbiosis^8–13^. Autophagy plays a vital role in the regenerative capacity of ISCs. Knockout (KO) of autophagy-related gene 5 (*Atg5*) in the intestinal epithelium of mice diminished the number of ISCs and reduced the intestinal organoid formation capacity *ex vivo*^14^. Moreover, ISCs isolated from the intestinal epithelium of *Atg7* or *Atg16L1* KO mice displayed decreased enteroid survival^15,16^. These observations support the requirement of autophagy for ISC function in mice. However, these results were obtained from a KO performed on all intestinal epithelial cell types, making it difficult to conclude an ISC-specific intrinsic role for autophagy.

*Drosophila melanogaster* has emerged as an invaluable tool to interrogate cell-specific roles of genes in intestinal functions^17^, given the multitude of genetic tools and reporters available^18^. Notably, the signaling pathways regulating ISC proliferation and differentiation are conserved from *Drosophila* to mammals, with less redundancy in flies^19^. Recent studies have demonstrated that the long-term RNAi-mediated depletion of diverse autophagy genes in ISCs and their progenitor cells causes ISC loss via apoptosis through DNA damage accumulation^20^. This loss of ISCs is reminiscent of the phenotype observed in a pan-intestinal epithelium *Atg7* KO mouse model^15^. In contrast, short-term RNAi-mediated depletion of *Atg* genes in fly ISCs and their progenitor cells increases their proliferation through the hyperactivation of the epidermal growth factor receptor (EGFR) pathway^21^. Hence, autophagy limits adult ISC proliferation in younger individuals and prevents premature apoptotic death during aging.

Autophagy may also affect intestinal cell differentiation. *Atg16L1* KO mice presented fewer secretory Paneth cells among all intestinal epithelial cells^9,16^. Atg14 or FIP200 KO intestinal epithelium displayed shortened villi^22^. The *Atg101* KO in the *Drosophila* intestine exhibits fewer absorptive enterocytes^23^. Nevertheless, no direct link has been reported regarding the control of intestinal cell differentiation by autophagy, especially because autophagy can control both the stemness and differentiation of other adult stem cells^24,25^. High basal autophagy levels preserve mouse hematopoietic cell stemness^26^. Conditional *Atg7* KO in mouse hematopoietic stem cells and progenitors blocks differentiation into mature neutrophils^27^, whereas *Atg5* deletion in naïve CD4^+^ T cells favors commitment to T lymphocyte helper TH9 cells^28^. In flies, autophagy induction is required for cyst cell self-renewal, whereas autophagy represses testis cyst cell progenitor differentiation^29^. Since autophagy affects intestinal stem cell regenerative capacities, differentiation could be affected; however, the mechanisms controlled by autophagy involved in the ISC behavior remain poorly understood.

At the cellular level, autophagy is a highly conserved catabolic process that regulates the lysosomal degradation of intracellular components^30^. An array of Atg proteins orchestrates the formation of a double-membrane phagophore, resulting in its elongation and cargo incorporation, or by selectively surrounding the components interacting with an autophagy cargo adaptor, such as Ref(2)P (SQSTM1/p62 in mammals)^31–33^. Ultimately, the phagophore seals and matures to form an autophagosome, fusing with lysosomes to degrade and recycle the sequestered constituents^34^. Basal autophagic flux, the ratio of autophagosome formation to autolysosome degradation, can be either increased or repressed in response to environmental cues. For example, nutrient deprivation inhibits the mammalian target of rapamycin complex 1 (mTORC1), which, when active, represses autophagy^35^, leading to Atg1/ULK1 activation and phosphorylation of several autophagy regulators^36^. Additionally, the fusion step requires coordinated actions between core Atg regulators and auxiliary proteins such as soluble *N*-ethylmaleimide-sensitive factor attachment protein receptors (SNAREs), tethering complexes, and RAB GTPase proteins^37,38^. Previous work from our laboratory has demonstrated the requirement for the endosomal small GTPase Rab21 in autophagosome-lysosome fusion^39^ in flies and mammals^40,41^. Once activated by the guanine exchange factor set-binding factor (Sbf), Rab21 promotes the trafficking of vacuolar-associated membrane protein 7 (Vamp7) from early endosomes to lysosomes^39^. Vamp7 in flies (and VAMP8 in mammals) is associated with synaptosomal-associated protein 29 (SNAP29) and autophagosomal syntaxin 17 (Stx17) to induce autophagosome-lysosome membrane fusion^42^. VAMP8/Stx17/SNAP29 is one of the two SNARE complex combinations involved in autophagosome-lysosome fusion^43–46^. Rab21 and VAMP8 have known functions in the gut, with Rab21 affecting nutrient-absorptive enterocyte survival in *Drosophila*^41^, and VAMP8 regulating mucus secretion by goblet cells in mice^47^. However, their roles in ISCs and their link to autophagy have not been explored in this context.

Herein, we show Rab21 loss and autophagy inhibition, specifically in adult ISCs, favor enteroendocrine cell differentiation. Mechanistically, a reduction in autophagic flux increases the activation of signal transducer and activator of transcription 92E (Stat92E) by acting downstream of Hopscotch/Janus kinase (Hop/JAK), causing an increase in the enteroendocrine cell population. Our data suggest basal autophagy limits enteroendocrine cell differentiation by regulating Stat92E in adult ISCs.

## RESULTS

### Rab21 depletion in adult ISCs increases progenitor and mature enteroendocrine cell populations

In *Drosophila*, ISCs divide into stem cells and progenitor cells, named enteroendocrine progenitor cells (EEPs) or enteroblasts, which differentiate into enteroendocrine cells or enterocytes, respectively^48,49^ (Figure 1A). Data from a genome-wide RNAi screen targeting intestinal stem and progenitor cells using the Esg-Gal4^ts^ driver highlighted autophagic genes and Rab21 as regulators of intestinal homeostasis^50^.

**Figure 1:**
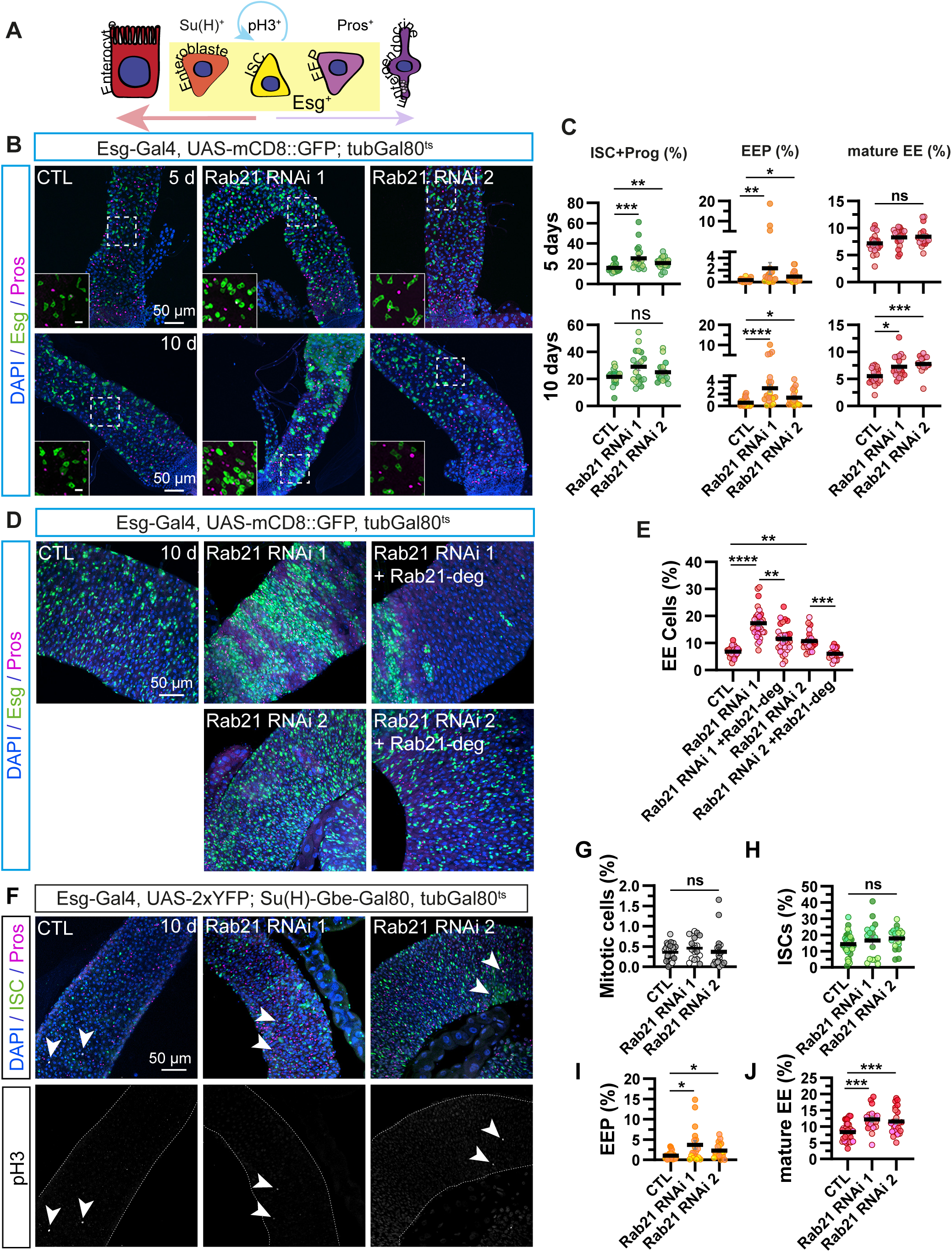
**Rab21 depletion in adult ISCs increases enteroendocrine cell populations**. (A) Schematic representation of intestinal cell lineages and cell-specific markers^70^: ISC, intestinal stem cell; EB, enteroblast; EEP, Enteroendocrine progenitor cell; Esg, Escargot; Su(H), suppressor of Hairless; Pros, Prospero; pH3, phospho-histone H3. (B–C) Adult *Drosophila* posterior midgut from Esg-Gal4, UAS-mCD8::GFP, tubGal80^ts^ driver expressing UAS-LacZ (CTL), Rab21 RNAi 1, or Rab21 RNAi 2 for five or ten days, in intestinal stem cells and progenitors. (B) Representative maximal projections. GFP labels Esg^+^ ISCs and progenitor cells (green). Prospero antibody marks enteroendocrine cells (magenta), and DAPI stains nuclei. Scale bar 50 µm. Magnification of GFP^+^, Pros^+^ enteroendocrine progenitor cells. Scale bar 10 μm. (C) Quantification of the percentage of GFP^+^ ISC and progenitor cells (left). GFP^+^, Pros^+^ enteroendocrine progenitor cells (middle). GFP^-^, Pros^+^ mature enteroendocrine cells (right). After 5 days (upper panel) or 10 days (lower panel) of RNAi expression. (D–E) Adult *Drosophila* posterior midgut from Esg-Gal4, UAS-mCD8::GFP, tubGal80^ts^ driver expressing UAS-LacZ (CTL) and co-expressing Rab21 RNAi 1 and 2 with UAS- LacZ or UAS-Rab21-deg for 10 days, in intestinal stem cells and progenitors. (D) Representative maximal projections. GFP labels Esg^+^ ISCs and progenitor cells (green). Prospero antibody marks enteroendocrine cells (magenta), and DAPI stains nuclei. Scale bar 50 µm. (E) Quantification of the percentage of total Pros^+^ enteroendocrine cells. (F–J) Adult *Drosophila* posterior midgut from Esg-Gal4, UAS-2xYFP; Su(H)-GBE-Gal80, tubGal80^ts^ driver expressing UAS-LacZ (CTL), Rab21 RNAi 1 or Rab21 RNAi 2 for 10 days in intestinal stem cells. YFP labels ISCs (green). Prospero antibody marks enteroendocrine cells (magenta), phospho-H3 labels the mitotic ISCs, and DAPI stains nuclei. Scale bar 50 µm. (F) Representative maximal projections. Quantification of the percentage of (G) pH3^+^ Mitotic cells. (H) YFP^+^, Pros^-^ ISC. (I) YFP^+^, Pros^+^ enteroendocrine progenitor cells. (J) YFP^-^, Pros^+^ mature enteroendocrine cells. Data information: N = three independent experiments from three independent crosses. Quantifications represent the mean ± SEM. Each dot represents an intestine. The Kruskal-Wallis test was used, followed by Dunn’s comparison tests. * p < 0.05, *** p < 0.001, **** p < 0.0001, ns non-significant p > 0.05. Also see Figure S1.

Specifically, the knockdown of Rab21, Atg2, and Atg6 induces hyperproliferation of ISCs and progenitor cells after seven days of depletion^50^. Given that autophagy inhibition affects ISCs differently depending on the knockdown duration^20,21^, we reinvestigated the requirement of Rab21 and autophagy in ISCs and progenitor cells in adult flies under homeostatic conditions. We used the targeting temporal and regional gene expression targeting (TARGET) system^51^, combined with the Esg-Gal4 driver, to deplete Rab21 using two previously validated independent RNAi constructs^39^. Intestinal cell populations were analyzed five or ten days after RNAi induction (Figure 1B and 1C). Green fluorescent protein (GFP) was expressed in Esg^+^ ISCs and progenitors (Figure 1A)^52^. Consistent with previous observations^50^, the depletion of Rab21 significantly augmented the percentage of GFP^+^ ISCs and progenitor cells after five days of RNAi induction (Figure 1B and 1C). However, the observed increase in GFP^+^ cells was no longer apparent ten days after Rab21 depletion (Figure 1B and 1C). Unexpectedly, the percentage of total enteroendocrine cells labeled with Prospero^53,54^ increased after five and ten days of knockdown (Figure S1A). We quantified mature enteroendocrine cells (Prospero^+^, GFP^-^) and EEPs (Prospero^+^ and GFP^+^)^52^. Five days after RNAi induction, Rab21 loss in ISCs and progenitor cells only induced more EEPs (Figure 1B and 1C). However, the depletion of Rab21 for ten days increased EEP and mature enteroendocrine cell populations (Figure 1B and 1C). We aimed to validate the specificity of Rab21 knockdown on the observed increase in the enteroendocrine cell population by expressing an untagged and degenerated Rab21 construct that was not targeted by Rab21 RNAis^41^. Significantly, the co-expression of degenerated Rab21 with Rab21 RNAis for ten days in ISC and progenitor cells rescued the increased proportion of Pros+ enteroendocrine cells (Figure 1D and 1E).

Since intestinal stem cells undergo asymmetric division to give rise to new ISCs and progenitor cells, we wondered whether the expansion in EEPs decreased the formation of enteroblasts. Using the Esg-Gal4^ts^ driver, we co-expressed the Notch reporter Su(H)-GFP^55^ with Rab21 RNAi to quantify Su(H)^+^ enteroblasts, and CFP^+^ ISCs and progenitor cells ^56^. Depletion of Rab21 for ten days did not affect the ratio of Su(H)^+^ enteroblasts to CFP^+^ ISCs and progenitor cells (Figure S1B). Therefore, an increase in enteroendocrine cells does not seem to be to the detriment of enteroblast formation. Taken together, our results show that Rab21 loss in ISCs and progenitor cells increases the number of EEPs and mature enteroendocrine cells without affecting the number of enteroblasts.

To explore the unique role of Rab21 in intestinal stem cells, we expressed Rab21 RNAis specifically in adult intestinal stem cells using an ISC-Gal4^ts^ driver line (Esg-Gal4, UAS-2xYFP; Su(H)-GBE-Gal80, tubGal80^ts^)^57^. ISC-specific Rab21 depletion for ten days did not affect ISC proliferation, as revealed by the mitotic marker phospho-histone 3 (pH3) (Figure 1F and G) and the percentage of total (YFP^+^, Prospero^-^) ISCs (Figure 1F and H). Significantly, the percentages of (YFP^+^, Prospero^+^) EEPs and (YFP^-^, Prospero^+^) enteroendocrine cells increased (Figure 1F, 1I and 1J). These results suggest Rab21 in adult intestinal stem cells modulates enteroendocrine cell proportion.

### Basal autophagy inhibition in ISCs augments the mature enteroendocrine cell population

Rab21 plays a role in autophagosome-lysosome fusion by regulating Vamp7 trafficking^39^. To confirm that Rab21 loss decreases autophagy in ISCs, we quantified the number of autophagy cargo adaptors Ref(2)P (SQSTM1/p62 in mammals) per ISCs (Figure 2A), as previously described in mammals and drosophila^31,58^. ISC-Gal4^ts^-driven knockdown of Rab21, Vamp7, and Stx17 led to Ref(2)P accumulation in YFP^+^ ISCs, whereas upon Sbf depletion, Ref(2)P dots increased to a lower degree (Figure 2A and 2B). These results are similar to those observed in various cell types following Rab21 depletion^39–41,59^. To assess whether the increase in enteroendocrine cells resulted from reduced autophagy upon Rab21 knockdown in adult ISCs, we artificially enhanced autophagy in Rab21- depleted ISCs via Atg1 overexpression or Tor inhibition. Atg1 overexpression and Tor inhibition are sufficient to increase autophagic flux^60–63^. Significantly, the increase in enteroendocrine cells caused by ISC-targeted Rab21 loss was rescued by Atg1 overexpression (Figure 2C and 2D). Tor inhibition was insufficient to completely restore the number of enteroendocrine cells. Differences in the observed effects may be caused by other non-autophagy-related functions of Tor. Additionally, since Rab21 also regulates the WASP and SCAR homologue (WASH) complex and the sorting of various cargo^64–67^, we tested whether the loss of WASH complex subunits would phenocopy Rab21 loss. Further supporting a direct link among Rab21, autophagy, and enteroendocrine cell formation, except for CCDC53, the knockdown of ISCs Wash, Strump, or FAM21, which are subunits of the Wash complex, did not affect the number of enteroendocrine cells (Figure S2A).

**Figure 2:**
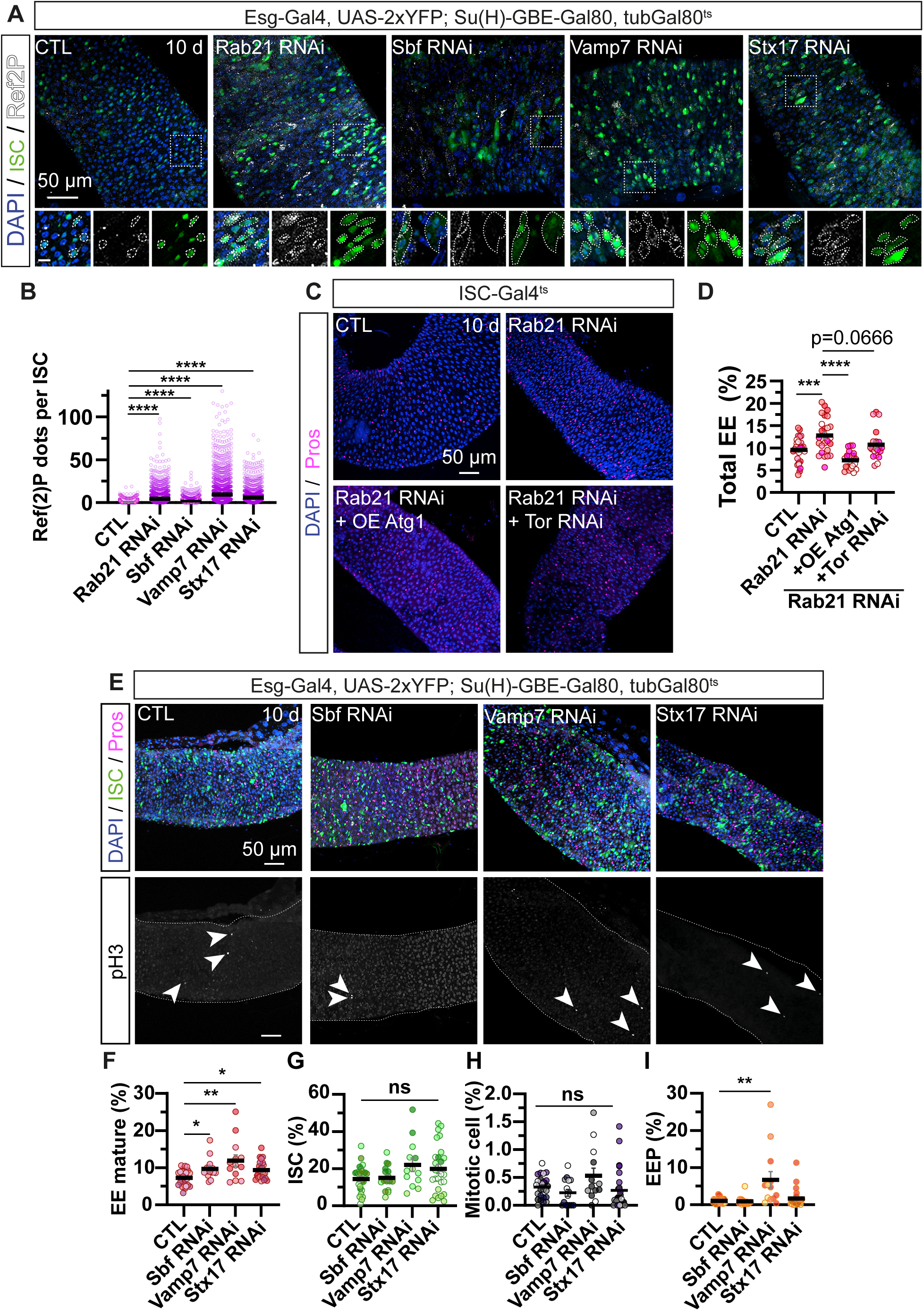
**Basal autophagy inhibition in intestinal stem cells augments the mature enteroendocrine cell population**. (A–B) Adult *Drosophila* posterior midgut from Esg-Gal4, UAS-2xYFP; Su(H)-GBE-Gal80, tubGal80^ts^ driver expressing UAS-LacZ (CTL), Rab21 RNAi 1, Sbf RNAi 2, Vamp7 RNAi, Stx17 RNAi for 10 days, in intestinal stem cells (A) Representative maximal projections. YFP labels ISCs (green). Ref(2)P antibody marks autophagosome cargos (white), and DAPI stains nuclei. Scale bar 50 μm. Magnification of Ref(2)P dots in YFP^+^ ISCs (dash lines). Scale bar 10 μm. (B) Quantification of Ref(2)P dots per YFP^+^ ISC. (C–D) Adult *Drosophila* posterior midgut from Esg-Gal4, UAS-2xYFP; Su(H)-GBE-Gal80, tubGal80^ts^ driver expressing UAS-LacZ (CTL) and co-expressing Rab21 RNAi 2 with UAS-LacZ or UAS-Atg1 or Tor RNAi for 10 days, in intestinal stem cells. (C) Representative maximal projections. Prospero antibody marks enteroendocrine cells (magenta), and DAPI stains nuclei. Scale bar 50 μm. (D) Quantification of the percentage of total Pros^+^ enteroendocrine cells. (E–I) Adult *Drosophila* posterior midgut from Esg-Gal4, UAS-2xYFP; Su(H)-GBE-Gal80, tubGal80^ts^ driver expressing UAS-LacZ (CTL), Rab21 RNAi 1, Sbf RNAi 2, Vamp7 RNAi, Stx17 RNAi for 10 days, in intestinal stem cells. (E) Representative maximal projections. YFP labels ISCs (green). Prospero antibody marks enteroendocrine cells (magenta), phospho-H3 labels the mitotic ISCs, and DAPI stains nuclei. Scale bar 50 μm. Arrowheads indicate pH3^+^ cells. Quantification of the percentage of (F) YFP^-^, Pros^+^ mature enteroendocrine cells. (G) YFP^+^, Pros^-^ ISC. (H) pH3^+^ mitotic cells. (I) YFP^+^, Pros^+^ enteroendocrine progenitor cells. Data information: N = three independent experiments from three independent crosses. Quantifications represent the mean ± SEM. Each dot represents an intestine. (B) Each dot represents Ref(2)P puncta per cell. The Kruskal-Wallis test was used, followed by Dunn’s comparison tests. * p < 0.05, *** p < 0.001, **** p < 0.0001, ns non-significant p > 0.05. Also see Figure S2 and S3.

Knockdown of Sbf, Vamp7, or Stx17 in ISCs for ten days using ISC-Gal4^ts^ phenocopied the increase in the (Pros^+^, YFP^-^) mature enteroendocrine cell population (Figure 2E and 2F) observed in Rab21-depleted intestines. Moreover, the increase in the total number of enteroendocrine cells upon Sbf depletion was confirmed using different RNAi (Figure S2B). Similar to Rab21 loss in ISCs, depletion of these regulators did not affect YFP^+^, Pros^-^ ISC number (Figure 2E and 2G), or the percentage of mitotic cells (pH3; Figure 2E and 2H), whereas the effect on YFP^+^ and Pros^+^ EEP cells was more subtle (Figure 2E and 2I). Similar to the loss of Rab21 in ISCs and progenitors (Figure S1B), the knockdown of Sbf or Vamp7 did not affect the ratio of enteroblasts compared with the number of CFP^+^ ISCs and progenitors (Figure S3). Altogether, these results suggest that limiting autophagosome-lysosome fusion in adult ISCs increases the number of mature enteroendocrine cells.

### Under homeostatic conditions, inhibiting basal autophagy favors the differentiation into enteroendocrine cells

To investigate if dysfunctional autophagy affects the differentiation into enteroendocrine cells, we used the Esg-Gal4 ReDDM (Repressible Dual Differential stability cell Marker) lineage-tracing method^68^. To label ISC progeny, stable histone H2B fused with RFP was co-expressed with RNAis against autophagy-related genes. Differentiated daughter mature enteroendocrine cells were Prospero^+^, GFP^-^, and RFP^+^, while mature enterocytes were Prospero^-^, GFP^-^, and RFP^+^ (Figure 3A). The percentage of RFP-labelled mature enteroendocrine cells derived from knocked-down ISCs and progenitors significantly increased upon Rab21, Sbf and Vamp7 depletion (Figure 3B and 3E, arrows).

**Figure 3:**
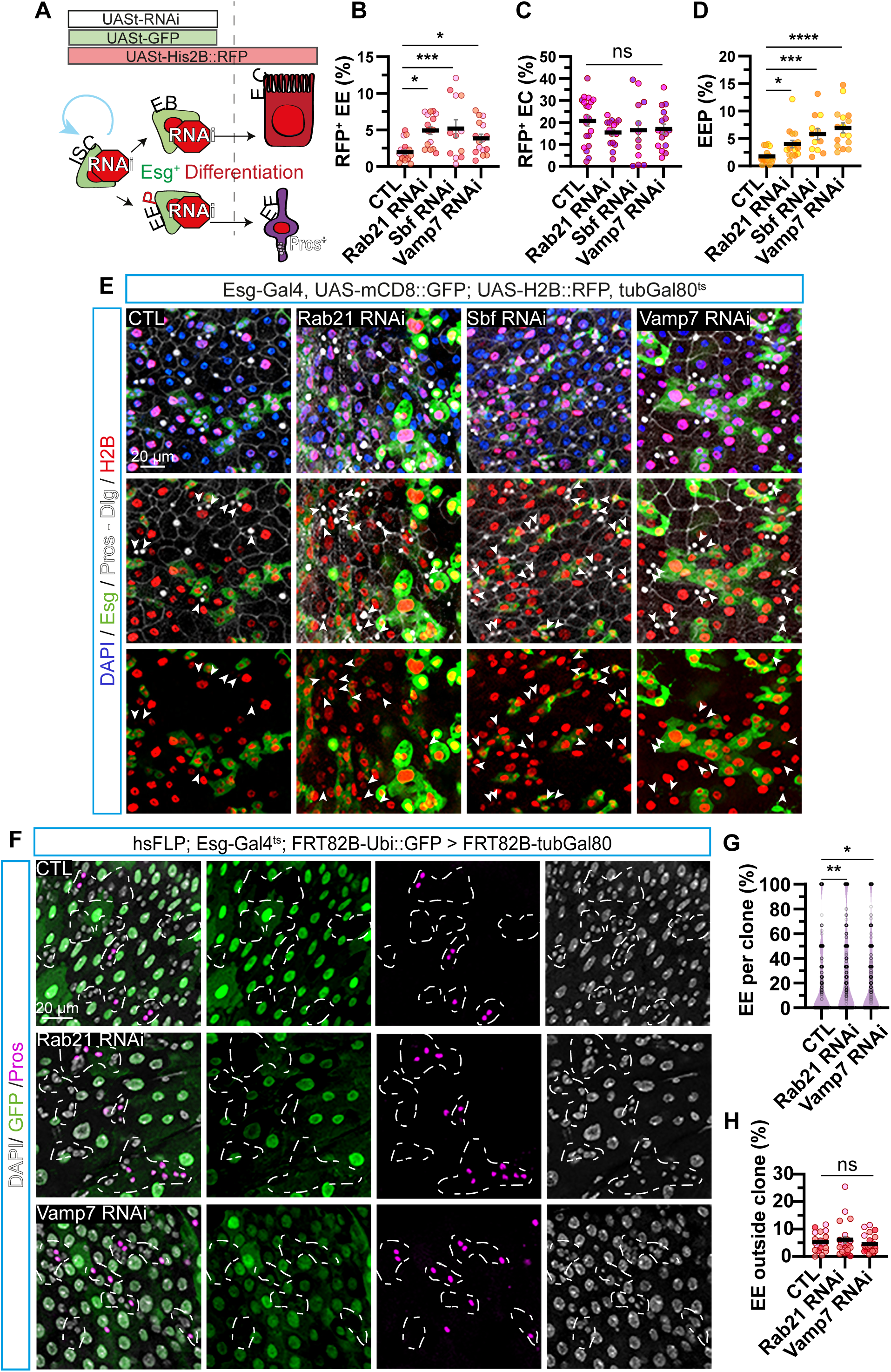
**Under homeostatic conditions, inhibiting basal autophagy favors the differentiation into enteroendocrine cells**. (A) Schematic representation of the ReDDM lineage-tracing method (B–E) Adult *Drosophila* posterior midgut from Esg-Gal4, UAS-mCD8::GFP; UAS- H2B::RFP, tubGal80^ts^ driver expressing UAS-LacZ (CTL), Rab21 RNAi 1, Sbf RNAi 1, Vamp7 RNAi for 10 days, in intestinal stem cells and progenitors. Quantification of the percentage of (B) (GFP^-^, RFP^+^ Pros^+^) enteroendocrine cells differentiated from depleted ISCs and progenitors. (C) (GFP^-^, RFP^+^ Pros^-^) enterocytes differentiated from depleted ISCs and progenitors. (D) (GFP^+^, RFP^+^, Pros^+^) enteroendocrine progenitor cells. (E) Representative maximal projections from the top to the lumen of the intestine. GFP labels Esg^+^ ISCs and progenitors (green). Pros and Dlg (Discs large) antibodies mark enteroendocrine cells and cell plasma membranes, respectively (white), H2B labels cell lineages from depleted ISCs and progenitors (red), and DAPI stains nuclei, arrowheads show RFP^+^ Pros^+^ enteroendocrine cells. Scale bar 20 µm. (F–H) MARCM from hsFlp, Esg-Gal4, tubGal80^ts^; FRT82B-UbiGFP cross with FRT82B (CTL), Rab21 RNAi 2; FRT82B-tubGal80, Vamp7 RNAi; FRT82B-tubGal80. Following the heat-shock recombination, RNAis were expressed for 10 days in intestinal stem cells and progenitors in the recombined clones. (F) Representative maximal projections from the top to the lumen of *Drosophila* posterior midgut. GFP^-^ labels clonal progenies delimited by dash lines. GFP^+^ labels non-recombined cells (green). Pros antibody marks enteroendocrine cells (magenta), and DAPI stains nuclei. Scale bar 20 µm. Quantification of the percentage of (G) Pros^+^ enteroendocrine cells per GFP^-^ clones. (H) GFP^+^, Pros^+^ enteroendocrine cells outside clones. Data information: (A–E) N = three independent experiments from three independent crosses; (F–H) N = three independent experiments from three and two independent crosses. Quantifications represent the mean ± SEM. (C, D, E, and G) Each dot represents an intestine. (G) Each dot represents a clone. The Kruskal-Wallis test was used, followed by Dunn’s comparison tests. * p < 0.05, *** p < 0.001, **** p < 0.0001, ns non-significant p > 0.05. Also see Figure S3.

Simultaneously, the number of enterocytes remained close to that of the controls (Figure 3C and 3E). Moreover, the knockdown of Rab21, Sbf and Vamp7 correlated with many EEPs (GFP^+^, RFP^+,^ and Prospero^+^) (Figure 3D and 3E). Therefore, inhibition of autophagy seems to enhance enteroendocrine differentiation without affecting enterocyte differentiation.

To further investigate the cell-autonomous effect of Rab21 and Vamp7 loss on enteroendocrine cell differentiation, we used the MARCM system^69^, which allows the labeling of single ISC daughter cell lineages and characterization of cell-autonomous and non-autonomous effects. By combining the Esg-Gal4 driver with the FRT82B-tubGal80 MARCM system, RNAi expression was restricted to ISCs and progenitors in GFP- generated clones. After ten days of RNAi induction, the loss of Rab21 or Vamp7 in ISC clones (represented by the dashed lines, Figure 3F) increased the percentage of enteroendocrine cells per clone (Figure 3F and 3G) without affecting their number outside the clones (Figure 3H), compared with controls. This suggests a cell-autonomous increase in autophagy-deficient ISC specification in the enteroendocrine cells. The ReDDM and MARCM lineage-tracing methods collectively suggest that basal autophagy inhibition, mediated by Rab21 or Vamp7 loss, autonomously promotes ISC and progenitor differentiation into enteroendocrine cells.

### Rab21 affects enteroendocrine differentiation by activating Stat92E

We aimed to characterize the interplay between autophagy inhibition and pathways affecting enteroendocrine cell differentiation. Under homeostatic conditions, Notch, JAK-STAT, EGFR, and Hippo are conserved signaling pathways that regulate stem cell behavior^70^ (Figure 4A). We postulated three models in which Rab21 loss in ISCs increased the percentage of enteroendocrine cells in the intestine. Rab21 loss could directly promote the differentiation of enteroendocrine cells. Alternatively, the higher number of enteroendocrine cells caused by Rab21 depletion may indirectly result from decreased enterocyte differentiation or increased ISC proliferation^21^. To test these hypotheses, we performed targeted screening of known modulators of these processes in Rab21-depleted ISCs and progenitor cells (Esg-Gal4^ts^). Briefly, enteroendocrine cell differentiation is inhibited by blocking the JAK-STAT pathway (Dome, Stat92E) or using a transcriptional inducer of enteroendocrine differentiation (Scute)^49,71,72^. We promoted enterocyte differentiation by overexpressing Delta or Sox21a^73–75^. Finally, we blocked ISC proliferation by knocking down EGFR, String (Stg), or Yorkie (Yki)^21,76^. Stat92E of the JAK-STAT pathway and the enteroendocrine cell fate inducer Scute were the most potent modifiers from this focused screen (Figure 4B and 4C). Overexpression of Delta or Sox21a to induce enterocyte differentiation did not rescue the increased proportion of EE from Rab21 loss (Figure 4B and 4C). This suggests that the higher number of enteroendocrine cells observed during Rab21 ISCs depletion was not an indirect consequence of fewer enterocytes. Delta overexpression did not modify Rab21 depletion, indicating that this effect was independent of the Notch pathway (Figure 4B and 4C). This was also supported by the fact that no difference in Su(H)-GFP intensity was observed in Rab21 knockdown intestines (Fig. S1B). Finally, the inhibition of ISC proliferation did not restore the number of enteroendocrine cells, indicating that this increase was not due to ISC accumulation. This screening of cell fate modulators suggests an interplay between Rab21 and the JAK-STAT pathway to regulate enteroendocrine cell differentiation.

**Figure 4:**
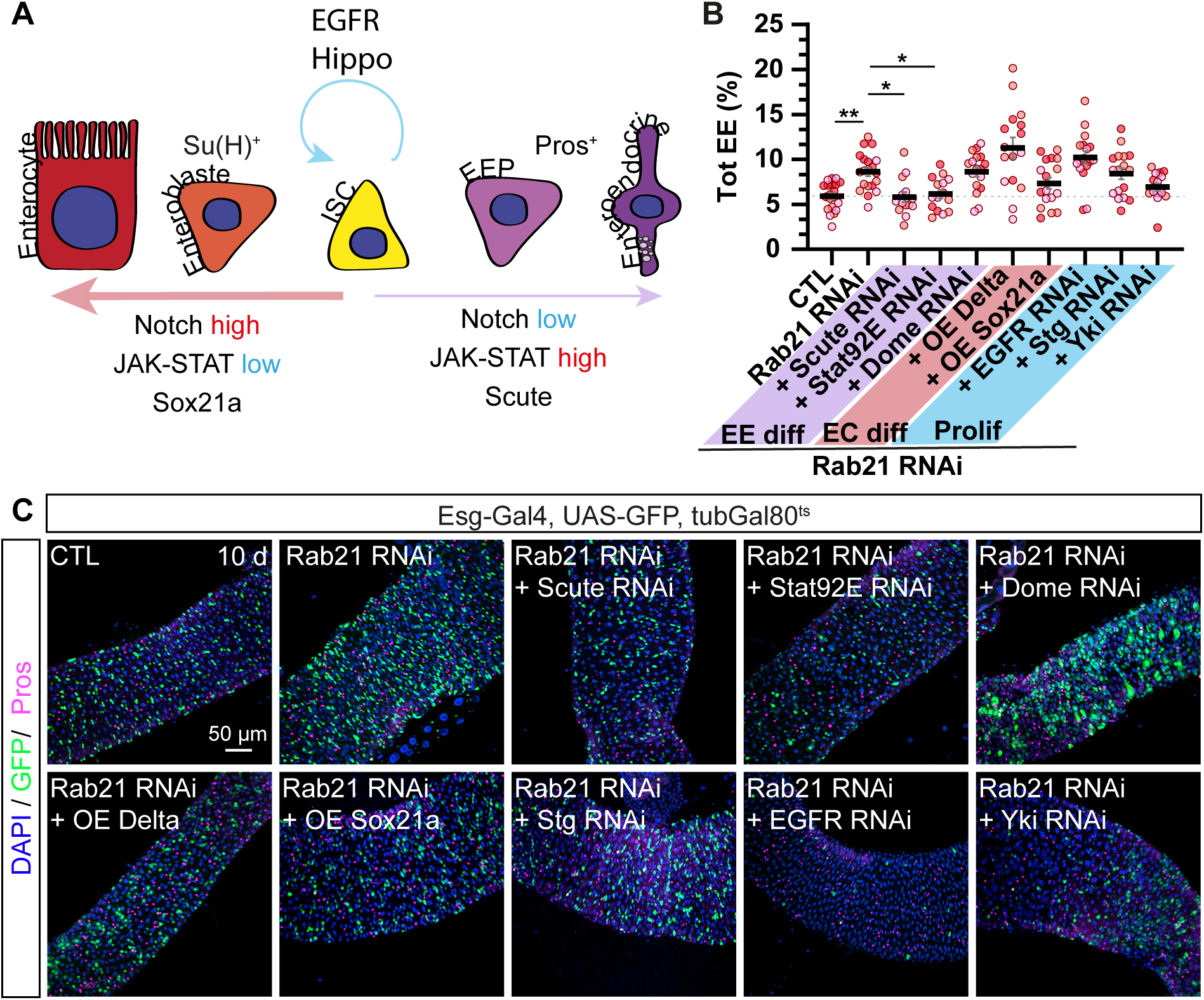
Rescue experiments with cell fate modulators identify Stat92E as a Rab21 modifier for enteroendocrine cell differentiation. (A) Schematic representation of signaling pathways regulating intestinal cell lineages: ISC, intestinal stem cell; EB, enteroblast; EEP, Enteroendocrine progenitor cell; Su(H), suppressor of Hairless; Pros, Prospero^70^. (B–C) Adult *Drosophila* posterior midgut from Esg-Gal4, UAS-GFP, TubGal80^ts^ driver expressing UAS-LacZ (CTL) or co-expressing Rab21 RNAi 2 with UAS-LacZ or with cell fate modulators for 10 days in intestinal stem cells and progenitors. (B) Quantification of the percentage of total Pros^+^ enteroendocrine cells. (C) Representative maximal projections. GFP labels ISCs and progenitor cells (green). Prospero antibody marks enteroendocrine cells (magenta), and DAPI stains nuclei. Scale bar 50 µm Data information: N = three independent experiments from three independent crosses. Quantifications represent the mean ± SEM. Each dot represents an intestine. The statistical tests used were one-way ANOVA followed by Dunnett’s comparison tests. * p < 0.05, *** p < 0.001, **** p < 0.0001, ns non-significant p > 0.05.

Therefore, we investigated the cell-autonomous effects of Rab21 in ISCs to understand how Rab21 genetically interacts with the JAK-STAT pathway. The JAK-STAT pathway (Figure 5A) is activated by the interaction between the membrane receptor Dome (Domeless) and secreted ligands Upd1-3 (Unpaired1-3), leading to Hop kinase autophosphorylation^77^. Hop (JAK ortholog) phosphorylates Stat92E, which dimerizes and translocates to the nucleus to induce target gene expression, including the suppressor of the cytokine signaling family, to repress the pathway^77^. Three Upds activate the JAK-STAT pathway, with Upd1 and Upd2 secreted by ISCs and progenitor cells, and Upd3 secreted by enteroblasts and enterocytes^77,78^. Since our focused screen targeted ISCs and progenitor cells (Esg-Gal4^ts^), we used the ISC-Gal4^ts^ driver and performed epistasis experiments, specifically in ISCs, to assess which JAK-STAT component(s) rescued Rab21 loss. Rab21 was co-depleted in ISCs along with Hop, Stat92E, Upd1, and Upd2. Depletion of Upd1 and Upd2 did not restore the increased enteroendocrine cell percentage (Figure 5B and 5D), which is in agreement with the lack of rescue by Dome RNAi in the original screen (Figure 4). Only Stat92E RNAi reduced the number of enteroendocrine cells, suggesting that Rab21 loss acts downstream of Hop to regulate enteroendocrine differentiation (Figure 5B, 5C, and 5D). Because heightened JAK-STAT signaling induces excessive enteroendocrine cell differentiation^72^, we wondered whether Rab21 depletion in ISCs could increase JAK-STAT activity in these cells. We monitored Stat92E activation with the 10XSTAT92E-GFP reporter^79^. Single-cell GFP relative fluorescence intensity measurements revealed a statistically significant increase in Rab21-depleted ISCs compared with control ISCs (Figure 5E and 5F). Taken together, our data show that by acting downstream of Hop/JAK, Rab21 inhibition in ISCs enhances Stat92E activity, thereby increasing enteroendocrine cell differentiation.

**Figure 5:**
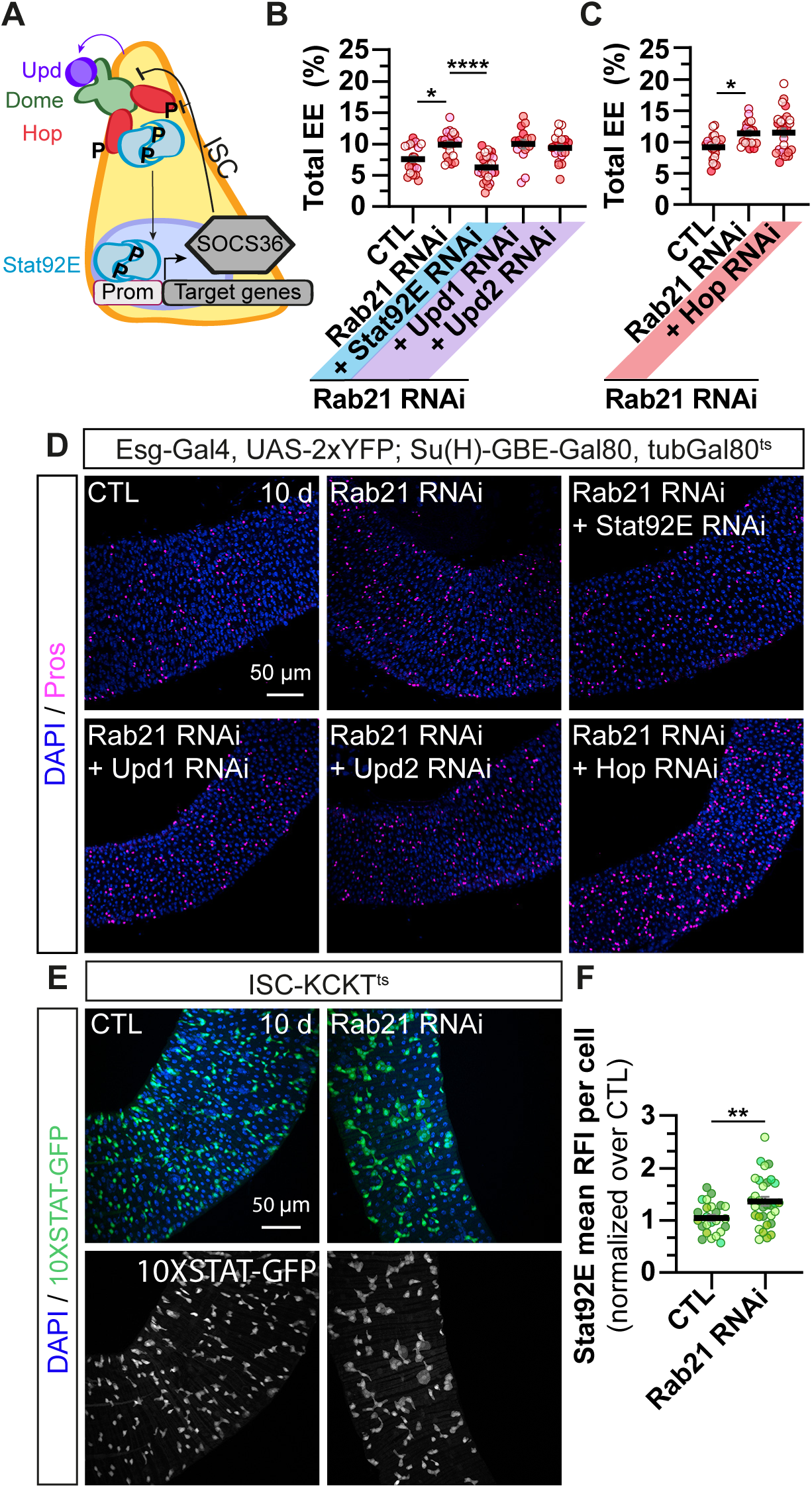
**Rab21 affects enteroendocrine differentiation by activating Stat92E downstream of Hop**. (A) Schematic representation of the JAK-STAT pathway^77^. ISC, intestinal stem cell; Upd, unpaired; Hop, Hopscotch; Dome, Domeless; Prom, Promoter; P, phosphorylation, SOCS36E, Supressor of cytokine signaling 36E. (B–D) Adult *Drosophila* posterior midgut from Esg-Gal4, UAS-2xYFP; Su(H)-GBE-Gal80, tubGal80^ts^ driver expressing UAS-GFP (CTL) and co-expressing Rab21 RNAi 2 with UAS-LacZ or Stat92E RNAi, Upd1 RNAi, Upd2 RNAi or Hop RNAi for 10 days, in intestinal stem cells. (B–C) Quantification of the percentage of total Pros^+^ enteroendocrine cells. (D) Representative maximal projections. Prospero antibody marks enteroendocrine cells (magenta), and DAPI stains nuclei. Scale bar 50 µm. (E–F) Adult *Drosophila* posterior midgut from ISC-KCKT^TS^ driver co-expressing 10XSTAT- GFP reporter with UAS-LacZ (CTL) or Rab21 RNAi 1. (E) Representative maximal projections. 10XSTAT-GFP labels cells with active Stat92E (green and white), and DAPI stains the nuclei. Scale bar 50 µm. (F) Quantification of 10XSTAT-GFP mean relative fluorescence intensity (RFI) per cell per intestine normalized to the control. Data information: N = four independent experiments from four independent crosses. Quantifications represent the mean ± SEM. Each dot represents the intestine. Statistical tests used were as follows: (B) Kruskal-Wallis test followed by Dunn’s comparison test, (C) One-way ANOVA test followed by Dunnett’s comparison test, and (D) unpaired t-test. * p < 0.05, *** p < 0.001, **** p < 0.0001, ns non-significant p > 0.05.

### Autophagy inhibition increases the enteroendocrine cell population through Stat92E activation

Our results suggest that inhibiting autophagy regulators involved in autophagosome-lysosome fusion favors enteroendocrine differentiation. To extend these findings, we tested whether disrupting autophagosome formation or elongation would phenocopy Rab21, Sbf, Vamp7, and Stx17 loss by increasing the number of enteroendocrine cells and Stat92E activity. Using the ISC-Gal4^ts^ driver, RNAi against Atg1 and Atg17, which control phagophore formation, and Atg5, Atg7, Atg8, and Atg16, which control autophagosome elongation, were specifically expressed in ISCs (Figure 6A and B). Since the 10XSTAT92E-GFP reporter labels ISCs and progenitors^75^, we validated RNAi efficiency on autophagy by quantifying the number of Ref(2)P dots in GFP-positive cells. The inhibition of all Atg genes increased the number of Ref(2)P dots (Figure S4A). Depletion for ten days of all the tested Atg genes increased the percentage of enteroendocrine cells and the activity of Stat92E, except for Atg16 (Figure 6A-6C). Atg16 genetically interacts in an autophagy-independent manner with the Slit-Robo2 pathway involved in enteroendocrine cell differentiation^80^. This may explain the increased number of enteroendocrine cells observed following Atg16 knockdown.

**Figure 6:**
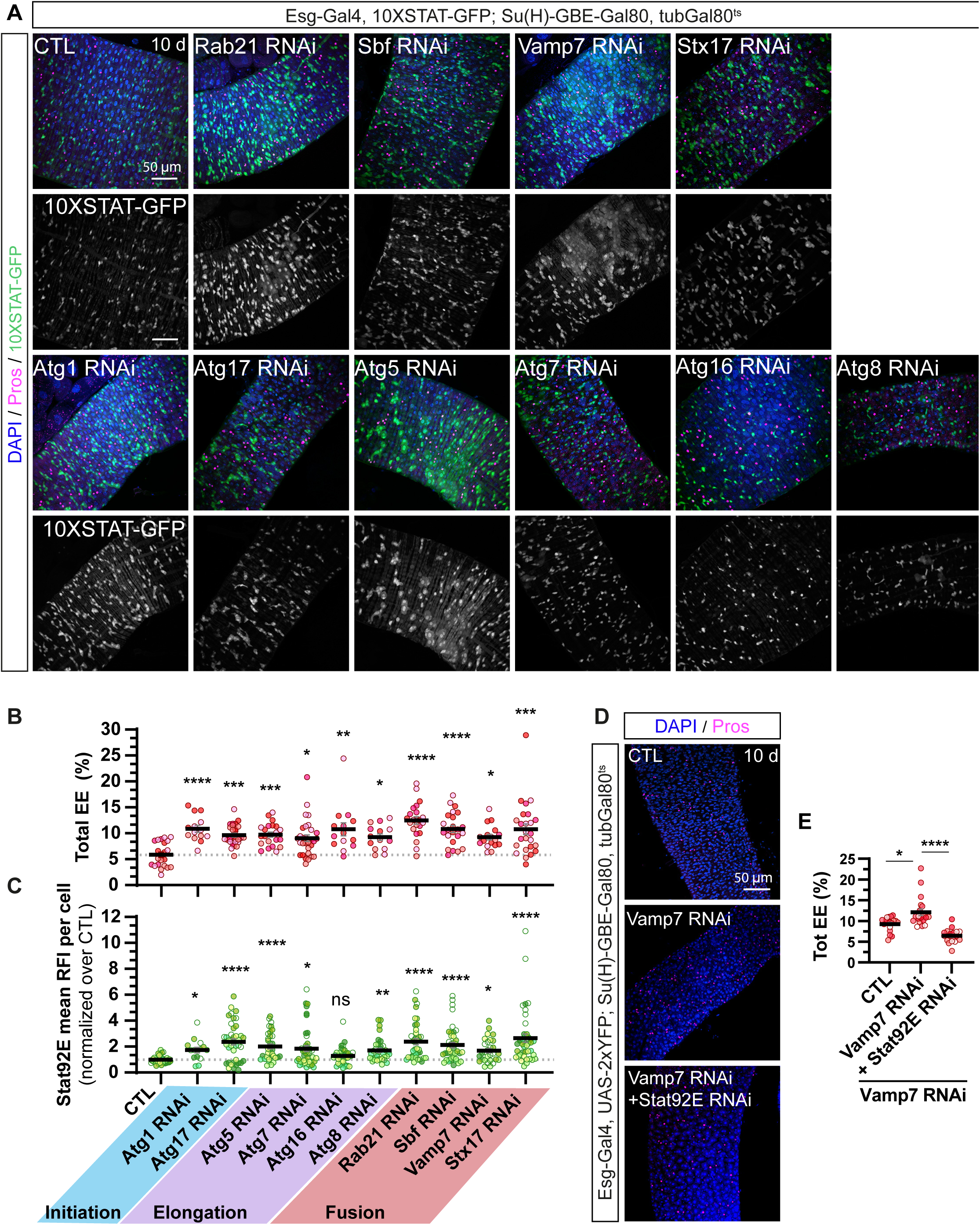
Disrupting autophagy in intestinal stem cells increases the enteroendocrine cell population through Stat92E activation. (A–C) Adult *Drosophila* posterior midgut from Esg-Gal4, 10XSTAT-GFP; Su(H)-GBE- Gal80, tubGal80^ts^ driver expressing UAS-mCD8:RFP (CTL) or RNAis against core Atg genes involved at different stages of autophagy for 10 days, in intestinal stem cells. (A) Representative maximal projections. 10XSTAT-GFP labels cells with active Stat92E (green and white). Prospero antibody marks enteroendocrine cells (magenta), and DAPI stains nuclei. Scale bar 50 µm. (B) Quantification of the percentage of total Pros^+^ enteroendocrine cells, and (C) mean per intestine of 10XSTAT-GFP RFI per cell normalized to the control, RFI (relative fluorescence intensity). (D–E) Adult *Drosophila* posterior midgut from Esg-Gal4, UAS-2xYFP; Su(H)-GBE-Gal80, tubGal80^ts^ driver expressing UAS-GFP (CTL) and co-expressing Vamp7 RNAi with UAS- GFP or Stat92E RNAi for 10 days, in intestinal stem cells. (D) Representative maximal projections. Prospero antibody marks enteroendocrine cells (magenta), and DAPI stains nuclei. Scale bar 50 µm. (E) Quantification of the percentage of total Pros^+^ enteroendocrine cells. Data information: N = 4 independent experiments from four independent crosses. Quantifications represent the mean ± SEM. Each dot represents an intestine. The Kruskal-Wallis test was used, followed by Dunn’s comparison tests. * p < 0.05, *** p < 0.001, **** p < 0.0001, ns non-significant p > 0.05. See also figure S4

In adult ISCs, constitutive activation of JAK-STAT for ten days (positive control) drastically increased both Stat92E activity and the number of enteroendocrine cells. (Figure S4B). This suggests that autophagy moderates the activity of Stat92E to limit enteroendocrine cell fate. To test this possibility directly, we inhibited Vamp7 along with Stat92E, specifically in ISCs, and quantified the percentage of enteroendocrine cells. Similar to the Rab21 loss, Stat92E depletion rescued the increase in the proportion of enteroendocrine cells caused by Vamp7 depletion (Figure 6D and 6E). From these experiments, we conclude that autophagy regulates Stat92E activation to affect enteroendocrine cell differentiation, at basal state.

## Discussion

Dysfunctional autophagy is associated with IBD, which causes chronic inflammation to impair the composition and function of the intestinal epithelium^4,13,81,83^. Although proper intestinal regeneration requires a balance between ISC proliferation and differentiation^3^, few studies have investigated the role of autophagy in ISCs. In addition to its essential role in mitigating stressors and its requirement for the degradation of proteins and organelles^85^, autophagy is emerging as a regulator of cell fate in various types of adult stem cells^24,25,86^. By targeting multiple autophagy regulators in ISCs and combining lineage-tracing experiments with a targeted screen, we demonstrated that autophagy inhibition increased Stat92E activity, causing an increase in enteroendocrine cells in the adult fly intestine. Our data provide the first evidence of the role of basal autophagy in the differentiation of enteroendocrine cells in the *Drosophila* adult intestine.

### In adult ISCs, basal autophagy cell autonomously limits enteroendocrine cell differentiation without affecting enterocytes

In intestinal stem cells, a ten-day depletion of core autophagy genes (Atg1, Atg5, Atg7, Atg8/LC3, and Atg17/FIP200), as well as auxiliary regulators (Sbf, Rab21, Vamp7, and Stx17), increased the percentage of enteroendocrine cells (Figures 1, 2, and 6). Moreover, lineage-tracing analyses confirmed the ISC cell-autonomous effect of autophagy inhibition on enteroendocrine cell differentiation without affecting enterocytes (Figure 3). Notch is a master regulator of intestinal cell fate and is conserved in mammals. High Notch activity commits ISC toward enterocytes. In contrast, Notch inhibition induces a complete loss of enteroblast formation, leading to the accumulation of ISC and enteroendocrine cells^73,82,84^. In addition, a subclass of enteroendocrine cells expressing tachykinin hormonal peptides requires Notch activation for differentiation^48,49,87^. A recent transcriptomic analysis indicated that this binary choice between enterocyte or enteroendocrine cell fate occurs in ISCs or enteroendocrine progenitor cells rather than in enteroblasts, as suggested before^48^. Genetic manipulation of the Notch pathway and proper expression of the Notch reporter (Su(H)-GFP) support the idea that autophagy might limit enteroendocrine cell differentiation independently of the Notch pathway.

### Under homeostatic conditions, in adult ISCs, autophagy limits Stat92E activity to modulate enteroendocrine cell differentiation

Under physiological conditions, opposingly to Notch, the hyperactivation of JAK-STAT in ISCs converts ISCs into enteroendocrine cells^72^. Using a 10XSTAT-GFP reporter, we showed that autophagy inhibition augmented Stat92E activity (Figures 5 and 6). Our previous study demonstrated that the loss of Rab21 in enterocytes increased the Yki-induced expression of Upd3. Secreted Upd3 upregulates JAK/STAT signaling and increases the number of enteroendocrine cells^41^. Herein, the depletion of the receptor Dome, and Upd1 and Upd2 ligands, and Yki in adult ISCs failed to restore the number of enteroendocrine cells in the intestine (Figures 4 and 5). These results suggest that autophagy acts downstream of Hop to restrict Stat92E transcriptional activity and enteroendocrine differentiation. However, the knockdown of Atg16 phenocopied the increase in the enteroendocrine cell population but not Stat92E activity. A previous study has reported that in adult *Drosophila*, independent of its role in autophagy, Atg16 limits enteroendocrine cell differentiation by genetically interacting with the Slit-Robo2 pathway^80^. Similar to our data, JAK-STAT signaling was not affected in organoids derived from intestinal crypts of mice KO for *Atgl16L1*^88^, whereas mouse organoids derived from a pan-intestinal epithelium *VAMP8* KO (mammalian ortholog of Vamp7) exhibited an increase in STAT3 activation^47^. This suggests that the activation of STAT transcription factor by autophagy in ISCs may be conserved in mammals. However, further studies are required to understand how autophagy controls Stat92E activity in ISCs. One hypothesis is that autophagy might directly degrade Stat92E. It has been reported that STAT5 can be degraded under basal conditions in T-cell lymphocytes^89^, as well as in mesenchymal cells to limit osteoblast differentiation^90^. Interestingly, during infection, orbiviruses induce autophagic degradation of STAT2 to replicate in cells^91^. Since viruses and other pathogens often hijack existing pathways to survive^92^, the role of autophagy in controlling STAT levels is plausible. Further supporting this hypothesis, STAT1, 2, and 4 contain LC3- interacting motifs^93^. Stat92E also contains an Atg8-interacting motif related to the LC3- interacting motif WxxL in mammals^94^. Through this sequence, Stat92E can be recruited to autophagosomes for degradation^95^.

Another hypothesis is that autophagy might degrade positive regulators of Stat92E activity. Phosphorylated STAT proteins can interact with transcriptional co-activators such as EP300/CREB-binding protein (p300/CBP), Nuclear Factor-KAPPA B (NF-κB), or PU.1/SPI-1^96–99^. To enhance STAT6 activity, p300/CBP is recruited by p100^100^. Autophagic degradation of p100 induces a non-canonical activation of NF-κB^101^. In contrast, in hematopoietic progenitor cells, autophagy degrades PU.1/Spi-1 transcription factor to repress Th9 lymphocyte differentiation^28^. Based on these studies, it would be interesting to investigate whether autophagy degrades or inhibits Nejire, Relish, or Ets98B, which are the Drosophila orthologs of p300/CBP, p100, and PU.1/SPI-1, respectively, to restrict Stat92E activity. JAK-STAT is required for cell differentiation, as STAT loss-of-function mutants show decreased enteroendocrine cell and enterocyte differentiation^72,75^. However, this is independent of Upd activation because the null Upd mutant *Df(1)os1A* does not affect ISC proliferation or differentiation^72^. This suggests the essential regulation of the JAK-STAT pathway under basal conditions. Our data revealed the intrinsic activation of Stat92E in ISCs and its modulation by autophagy to regulate enteroendocrine cell differentiation.

### Autophagy participates in the control of stem cell behavior

Previous studies have reported differences in stem cell behavior in response to dysfunctional autophagy. A decrease in proliferation^20^ associated with a loss of ISC number^14,15,20^ has been reported in flies depleted of autophagy genes in ISCs and progenitors and in *Atg5* or *Atg7* KO mice in all epithelial cells. On the contrary, in other contexts, autophagy inhibition increased ISC proliferation^21,50^ both cell- and non-cell autonomously through EGFR and Yki activation, respectively^21^. Similar to these studies, the loss of Rab21 in ISCs and progenitor cells for five days increased their number. However, in ISCs, the knockdown of Rab21, Sbf, Vamp7, or Stx17 for ten days did not affect ISC proliferation, as shown by pH3 labeling (Figure 2). In line with these results, restricting ISC proliferation caused by the knockdown of EGFR, Stg, or Yki did not rescue the increase in the number of enteroendocrine cells (Figure 2). Thus, the increase in enteroendocrine cell differentiation was independent of ISC proliferation. Contrary to our data, previous MARCM analyses of Atg6, FiP200 KO, and SH3PX1 did not show any effect on intestinal cell differentiation^21^. However, these studies used the MARCM methodology, which would lead to knockdown or KO of the tested genes in ISCs, progenitors, but also in differentiated cells. Because autophagy plays an important role in enterocytes, its effect on enteroendocrine cells may have been overlooked. We observed an effect on enteroendocrine cells because we inhibited autophagy for ten days, specifically in ISC and progenitor cells. Similarly, a recent study demonstrated that the mTORC1 pathway suppressed enteroendocrine cell formation in response to nutrient adaptation^102^. However, this is region-specific and mainly regulated by the Delta-Notch pathway. Our study focused on the posterior midgut, which is sensitive to mTORC1-dependent cell-fate adaptation^102^. However, the authors did not assess the requirement for autophagy in the mTORC1-driven effect, but this new study^102^ might suggests a second independent mechanism by which autophagy modulation affects cell fate determination in the adult fly intestine. All these pieces of evidence suggest that the duration of autophagy inhibition can affect stem cell behavior differently.

### Dysfunctional enteroendocrine cells are associated with IBD

Our study established a role for autophagy in ISC fate decisions toward enteroendocrine cell differentiation in the unstressed intestinal epithelium. As pathogenic infections can increase JAK-STAT signaling to drive enteroendocrine cell differentiation in flies^103^, it will be of interest to test whether autophagy affects Stat92E activity following infection. Since infection-dependent activity requires ISC proliferation and depends on Dome and Hop^103^, autophagy could differentially regulate Stat92E activity based on the cellular context. Therefore, understanding how autophagy regulates the ISC fate in response to environmental changes is a significant new research direction for future studies. In mammals, STAT5 and STAT3 KOs in all intestinal epithelial cells are detrimental to intestinal crypt regeneration, whereas the addition of interleukins improves intestinal crypt regeneration without being necessary for organoid generation^104,105^. However, little is known about the role of STAT transcription factors in ISCs. Translating our results to mammals could help us understand the regulation and function of JAK-STAT in the autophagy-defective mammalian intestinal epithelium. Autophagy and JAK-STAT components, as well as Phox2B enteroendocrine cell regulator, have been identified as IBD susceptibility genes^106,107^. An increase in enteroendocrine cells has been reported in patients with Crohn’s disease^108^ as well as dysregulation of hormonal peptide secretion, which might correlate with dysfunctional transit in patients with IBD^109^. Similar to flies, infections in mice can lead to an increase in enteroendocrine cell subtypes^110^. Enteroendocrine cells regulate digestion and interact with immune cells and the stem cell niche^109^. Enteroendocrine cells can replace ablated Paneth cells, which are dysfunctional in IBD, to maintain the stem cell niche. Determining whether this interplay between autophagy, JAK-STAT, and enteroendocrine cells is conserved in the mammalian intestine could help define altered mechanisms affected in pathogenic conditions, such as IBDs, given the increased use of JAK inhibitors in patients with IBD^111^.

## MATERIAL AND METHODS

| Reagent or Resource | Source | Identifier |
| --- | --- | --- |
| Drosophila lines: Transgenes |  |  |
| UAS-Rab21 RNAi 1 | VDRC | 32941 |
| UAS-Rab21 RNAi 2 | VDRC | 109991 |
| UAS-Sbf RNAi 1 | VDRC | 22317 |
| UAS-Sbf RNAi 2 | Bloomington | 32419 |
| UAS-Stx17 RNAi | Bloomington | 25896 |
| UAS-Vamp7 RNAi | NIG | 1599R-1 |
| UAS-Atg1 RNAi | VDRC | 16133 |
| UAS-Atg5 RNAi | VDRC | 104461 |
| UAS-Atg7 RNAi | VDRC | 45558 |
| UAS-Atg8 RNAi | Bloomington | 34340 |
| UAS-Atg16 RNAi | Bloomington | 34358 |
| UAS-Atg17 RNAi | Bloomington | 36918 |
| UAS-Wash RNAi | VDRC | 39769 |
| UAS-Tor RNAi RNAi | Bloomington | 33951 |
| UAS-CCDC53 RNAi | VDRC | 28008 |
| UAS-Strump RNAi | Bloomington | 51906 |
| UAS-FAM21 RNAi | Bloomington | 50571 |
| UAS-VPS26 RNAi | Bloomington | 38937 |
| UAS-Upd1 RNAi | Bloomington | 28722 |
| UAS-Upd2 RNAi | Bloomington | 33988 |
| UAS-Dome RNAi | Bloomington | 32860 |
| UAS-Hop RNAi | Bloomington | 31319 |
| UAS-Stat92E RNAi | Bloomington | 33637 |
| UAS-Stg RNAi | Bloomington | 29556 |
| UAS-EGFR RNAi | Bloomington | 25781 |
| UAS-Scute RNAi | Bloomington | 26206 |
| UAS-ATG1 | Bloomington | 51655 |
| UAS-Delta-GFP | Bloomington | 8611 |
| UAS-Hop | Bloomington | 79033 |
| UAS-Sox21a | Bloomington | 68156 |
| UAS-LacZ | Bloomington | 1776 |
| UAS-GFP | Bloomington | 35786 |
| FRT82B | Bloomington | 5188 |
| UAS-mCD8:RFP | Bloomington | 27391 |
| 10XSTAT-GFP | Bloomington | 26197 |
| UAS-Rab21 RNAi 2; FRT82BGal80 |  | This Paper |
| UAS-Vamp7 RNAi; FRT82BGal80 |  | This Paper |
| UAS-Rab21 RNAi 1, 10XSTAT-GFP |  | Nassari et al,<br>2022 |
| UAS-LacZ, 10XSTAT-GFP |  | Nassari et al,<br>2022 |
| UAS-Rab21 degenerate |  | Nassari et al,<br>2022 |
| Drosophila lines: Drivers |  |  |
| Esg-Gal4, UAS-GFP, TubGal80<sup>ts</sup>/ CyO | Edgar B. |  |
| Esg-Gal4, UAS-mCD8:GFP/ CyO; Gal80<sup>ts</sup> | Biteau B. |  |
| Esg-Gal4, UAS-YFP/ CyO; Su(H)-GBE-Gal80,<br>tubGal80<sup>ts</sup>/ TM3, Sb | Bardin A. |  |
| Esg-Gal4, UAS-mCD8::GFP/ CyO; UAS-H2B::RFP,<br>tubGal80GAL80<sup>TS</sup>/ TM3,Ser | Dominguez M. |  |
| Esg-Gal4, UAS-CFP, Su(H)-GFP/ CyO, tubGal80<sup>ts</sup> /<br>TM3, Sb | O'Brien L. |  |
| hsp70-FLP; Esg-Gal4, tubGal80<sup>ts</sup> / CyO; FRT82B-Ubi::GFP | O'Brien L. |  |
| {gbe-GAL80}ZH-2A; {mira -KDRT>-dSTOP-KDRT>-<br>GAL4}attP40, P{tubP-GAL80[ts]}20; {CG10116-KD.PEST}attP2) | Sokol N. |  |
| Esg-Gal4, 10XSTAT-GFP/ CyO; GBE-Su(H)Gal80, tubGal80<sup>ts</sup>/ TM3,Ser | This Paper |  |
| Reagent or Resource | Source | Identifier |
| Antibodies |  |  |
| Anti-Prospero | DSHB | M1RA |
| Anti-phospho-histone H3 (Ser10) | Millipore | 06-570 |
| Anti-Ref(2)P | Abcam | ab178440 |
| Anti-discs large | DSHB | 4F3 |
| Goat anti-Mouse IgG (H+L) Secondary Antibody, Alexa Fluor® 647 conjugate | Thermo Fisher Scientific | A21236 |
| Goat anti-Rabbit IgG (H+L) Secondary Antibody, Alexa Fluor® 546 conjugate | Thermo Fisher Scientific | A11035 |
| Chemical reagents |  |  |
| Glycerol | Fisher Scientific | AAJ61059K2 |
| 20 % Formaldehyde Aqueous Solution (Paraformaldehyde Aqueous Solution) EM Grade | Electron microscopy science | 15713 |
| DAPI | New England Biolabs | 4083S |
| SlowFade™ Gold Antifade Mountant | Thermo Fisher Scientific | s36940 |
| SlowFade™ Gold Antifade Mountant with DAPI | Thermo Fisher Scientific | S36938 |
| Software |  |  |
| Image J | https://fiji.sc/ |  |
| CellProfiler | https://cellprofiler.org/releases |  |
| IMARIS | Oxford Instruments |  |
| Prism | GraphPad |  |
| Photoshop | Adobe |  |
| Illustrator | Adobe |  |

## Drosophila

The Drosophila stocks were conserved at room temperature (RT). Stocks and experimental flies were raised on a standard diet composed of 7 g/L agar, 60 g/L cornmeal, 60 g/L molasses, 23.5 g/L yeast extract, 4.5 mL/L Tegosept (BioShop), and 4 mL/L propionic acid.

## TARGET SYSTEM

Cell-specific Gal4 lines were combined with a ubiquitously expressed thermosensitive Gal80, referred to as Gal4^ts^, to repress Gal4 transcriptional activity at a permissive temperature (18–20°C). Experimental crosses were maintained at 20°C to inhibit GAL4 activity until eclosion. F1 female flies were collected 1–3 days after eclosion and shifted to 29°C to alleviate Gal80 repression for five or 10 days (see figure legends). Every two days, flies were flipped onto fresh food.

## MARCM

Hsp70-FLP; Esg-Gal4, tubGal80^ts^ / CyO; FRT82B-Ubi::GFP driver was crossed with FRT82B (CTL), Rab21 RNAi 2; FRT82B-tubGal80 or Vamp7 RNAi; FRT82B-tubGal80.

One- to three-day-old females were transferred to vials containing a buzz plug with water during heat shock. Heat shock induces the recombination of FRT sites, removing both GFP and Gal80 from one ISC daughter cell. Flies were incubated for 1 h in a water bath at 37°C and then shifted to 18°C for 2 h onto fresh food. The procedure was repeated on the same day. The flies were transferred to 29°C for ten days before dissection.

### Intestine dissection and immunocytochemistry

The intestines from adult female flies were dissected in phosphate-buffered saline (PBS), fixed with 4% paraformaldehyde for 2 h at RT, and transferred to PBS. To clean the lumen, the intestines were trimmed anteriorly and posteriorly to the posterior midgut, incubated in glycerol 50% in PBS for 20 min, and washed three times for 10 min with PBS-0.1%Triton at RT. After overnight incubation at 4°C with the primary antibodies diluted in PBS-0.1%Triton, the intestines were washed three times for 10 min at RT with PBS-0.1%Triton. Secondary antibody incubation was performed for at least 3 h at RT in PBS-0.1%Triton, followed by three times 10-minute incubations with PBS-0.1%Triton, and three times 10-minute incubations with PBS at RT. Nuclei were stained with DAPI (1:10000) diluted in PBS for 20 min and rinsed twice with PBS for 10 min at RT. The intestines were incubated for 10–20 min in 50% glycerol PBS before being mounted in SlowFade Gold Antifade Mountant with or without DAPI. The following primary and secondary antibodies were used: mouse anti-Prospero (DSHB, 1:1000), rabbit anti-pH3 (Millipore, 1:1000), rabbit anti-Ref(2)P (Abcam, 1:1000), mouse anti-discs large (1:250), Alexa anti-mouse 647 (Invitrogen, 1:1000), and Alexa anti-rabbit 546 (Invitrogen, 1:1000).

### Image acquisition

The posterior midgut R5 regions were imaged on a Zeiss LSM880 confocal microscope using either a 20× Plan-APOCHROMAT/0.8 numerical aperture (NA) or a 40× oil Plan-APOCHROMAT/1.4 NA objective. Z-stacks were performed from one side to the other of the midgut with 2 μm steps between images. For intensity measurement using the 10XSTAT-GFP reporter, z-stacks were acquired from one side until the lumen part with 1 μm steps between images. Representative images are maximal projections unless specified otherwise. The image levels were adjusted uniformly and linearly for all conditions within the same experiment using Photoshop software.

### Image analysis

All images were analyzed using an automated approach.

Z-stack images were analyzed in 3D using the IMARIS software to quantify intestinal cell populations. First, images were processed to diminish the background and to smooth the signal by applying a Gaussian filter of 0.6 μm and a background subtraction of 10 μm, 5 μm, and 20 μm for nuclei (DAPI or RFP^+^), Pros^+^ cells, and ISC or ESG cells, respectively. Second, the objects were detected using spot or surface modules. Spots identified DAPI-, Prospero^+^ cells, RFP^+^ cells, Esg-CFP^+^ cells, Su(H)^+^ cells, or Ref(2)P dots. Surfaces were created to identify ISC^+^ and Esg ^+^ cells expressing YFP or GFP, respectively. Third, the “Find spots close to surface” module was used with a 0.5 or 1 μm threshold to identify co-labeled ISC^+^ or Esg^+^ cells with Prospero, RFP, or DAPI spots. This module distinguishes “spots close to the surface” representing ISC^+^ or Esg^+^ cells from “spots far from the surface” representing ISC^-^ or Esg^-^ cells. The “Colocalized spots” module was used with a threshold of 5 μm to identify (Pros^+^, RFP^+^) cells or (RFP^+^, DAPI^+^) cells. Ref(2)P dots per ISC were quantified using the “split spot into surface” module.

Measurement of 10XSTAT-GFP intensity was performed using CellProfiler. Maximal projections were generated using ImageJ software. First, the background was reduced using “rescale intensity” and “reduce noise” modules. Second, 10XSTAT-GFP^+^ cells were delimited from the processed images using “Identify primary object.” Finally, the “Measure object intensity” module quantified the integrated GFP intensity from the unmodified images within the 10XSTAT-GFP^+^ cells primary object. For each experiment, the average intensity per cell per intestine was calculated and normalized to the average intensity of CTLs.

MARCM analysis was performed using CellProfiler. The background was removed using “rescale intensity,” “Enhance speckles feature,” “Gaussian Filter,” and/or “reduce noise” modules. DAPI, Pros+, and GFP+ were determined with the “identify primary objects” module. “Masked objects identified” (DAPI, GFP^-^), (Pros^+^, GFP^-^) cells, and (Pros^+^, GFP^+^) cells. “Overlay objects” and “Overlay outlines” modules were applied to create an image of the GFP channel merging the previously identified objects: GFP^+^ cell objects, (DAPI, GFP^-^) objects, and (Pros^+^, GFP^-^) objects. This generated image was used to manually draw GFP^-^ clones with the “identify objects manually” module. Finally, the number of DAPI or Pros+ cells per clone was calculated from “relate objects,” which assessed the child object (DAPI, GFP^-^) or (Pros^+^, GFP^-^) identified above to the parent object corresponding to the manually drawn clone.

### Statistical analyses

All experiments were performed at least three times from three independent crosses (two independent crosses were performed for MARCM analysis). Statistical analyses were performed using the GraphPad Prism software. Normality was first assessed, followed by a parametric or nonparametric test, depending on the outcome of the normality test. An unpaired t-test or Mann-Whitney test was performed if only two conditions were compared. For multiple comparisons, one-way analysis of variance (ANOVA) or Kruskal-Wallis tests were performed, followed by Dunnett’s or Dunn’s tests. P-values are represented as * p < 0.05, *** p < 0.001, **** p < 0.0001, and ns = non-significant p >0.05. Graphs representing data show means ± S.E.M.

## ACKNOWLEDGMENTS

We thank Bruce Edgar for kindly providing the transgenic fly lines, Esg-Gal4, UAS-GFP, and TubGal80ts/ CyO. We thank Alison Bardin for kindly providing the transgenic fly lines Esg-Gal4, UAS-YFP/ CyO, Su(H)-GBE-Gal80, and tubGal80ts/ TM3,Sb. We thank Benoit Biteau for kindly providing the transgenic fly line Esg-Gal4, UAS-mCD8:GFP/CyO; Gal80^ts^. We thank Maria Dominguez for kindly providing the transgenic fly line Esg-Gal4, UAS-mCD8::GFP/ CyO; UAS-H2B::RFP, tubGal80GAL80TS/ TM3,Ser. We thank Lucy O’Brien for kindly providing the transgenic fly lines Esg-Gal4, UAS-CFP, Su(H)-GFP/CyO, tubGal80ts / TM3,Sb, and hsp70-FLP; Esg-Gal4, tubGal80ts/ CyO; FRT82B-Ubi::GFP. We thank Nicolas Sokol for kindly providing us with the ISC-KCK-TS fly line. We thank Rajan Akhila for kindly providing the fly line UPD2-HA. We would like to thank Editage (www.editage.com) for English language editing. We thank the photonic microscopy platform used for confocal microscopy. We also thank all the members of the Jean Laboratory for providing relevant suggestions during this work.

## Author contributions

C.L-K. conceived, performed and analyzed all the data. C.L-K wrote the first version of the manuscript and prepared the various figures. S.N. performed and analyzed specific experiments. S.J. supervised, acquired funding and edited the manuscript for publication.

