## Supplemental Figures for "Autophagy inhibition in intestinal stem cells favors enteroendocrine cell differentiation through Stat92E activity"

### Supplemental information

Document S1. Figure S1-S4

### SUPPLEMENTARY FIGURE LEGENDS

**Figure S1: Rab21 depletion in intestinal stem cells and progenitor cells affects the total number of enteroendocrine cells without affecting enteroblasts.**

(A) Quantification of the percentage of total $Pros^+$ enteroendocrine cells related to Figure 1B.

(B) Adult *Drosophila* posterior midgut from Esg-Gal4, UAS-CFP, Su(H)-GBE-GFP, $tubGal80^{ts}$ driver expressing UAS-LacZ (CTL), RAB21 RNAi 1 or 2 for 10 days, in intestinal stem cells and progenitors. Representative maximal projections. CFP labels ISCs and progenitor cells (blue). GFP marks enteroblasts (EB). Scale bar 50 µm. The graph represents the quantification of the ratio of the number of GFP+ EB over the number of $CFP^+$ ISCs and progenitors.

Data information: N = three independent experiments from three independent crosses. Quantifications represent the mean ± SEM. Each dot represents an intestine. The statistical tests used were: (A) unpaired t-tests. (B) Kruskal-Wallis test followed by Dunn's comparison test. * $p < 0.05$, *** $p < 0.001$, **** $p < 0.0001$, ns non-significant $p > 0.05$.

**Figure S2: Targeting Sbf with a different RNAi phenocopy the increase in enteroendocrine cells, while the knockdown of most of the Wash complex subunits does not.**

(A) Adult *Drosophila* posterior midgut from Esg-Gal4, UAS-2xYFP; Su(H)-GBE-Gal80, $tubGal80^{ts}$ driver expressing UAS-LacZ (CTL), Rab21 RNAi 2 or RNAis against subunits of the Wash complex, Wash, CCDC53, FAM21 or Strump for 10 days in intestinal stem cells. Prospero antibody marks enteroendocrine cells (magenta), and DAPI stains nuclei. Representative maximal projections. Scale bar 50 µm. Graphs represent the quantification of the percentage of $Pros^+$ mature enteroendocrine cells.

(B) Adult *Drosophila* posterior midgut from Esg-Gal4, UAS-2xYFP; Su(H)-GBE-Gal80, $tubGal80^{ts}$ driver expressing UAS-LacZ (CTL), Sbf RNAi 2 for 10 days in intestinal stem cells. Prospero antibody marks enteroendocrine cells (magenta), and DAPI stains nuclei. Representative maximal projections. Scale bar 50 µm. The graph represents the quantification of the percentage of $Pros^+$ mature enteroendocrine cells.

Data information: N = three independent experiments from three independent crosses. Quantifications represent the mean ± SEM. Each dot represents an intestine. The Kruskal-Wallis test was used, followed by Dunn's comparison tests. * $p < 0.05$, *** $p < 0.001$, **** $p < 0.0001$, ns non-significant $p > 0.05$.

**Figure S3: Similarly to Rab21, Sbf or Vamp7 depletion in intestinal stem cells and progenitor cells does not affect the number of enteroblasts.**

Adult *Drosophila* posterior midgut from Esg-Gal4, UAS-CFP, Su(H)-GBE-GFP, tubGal80$^{ts}$ driver expressing UAS-LacZ (CTL), Sbf RNAi 1 or Vamp7 RNAi for 10 days, in intestinal stem cells and progenitors. Representative maximal projections. CFP labels ISCs and progenitor cells (blue). GFP marks enteroblasts (EB). Scale bar 20 μm. The graph represents the quantification of the ratio of the number of GFP+ EB over the number of CFP$^+$ ISCs and progenitors.

Data information: N = three independent experiments from three independent crosses. Quantifications represent the mean ± SEM. Each dot represents an intestine. The Kruskal-Wallis test was used, followed by Dunn's comparison tests. * $p < 0.05$, *** $p < 0.001$, **** $p < 0.0001$, ns non-significant $p > 0.05$.

**Figure S4: Intrinsic overactivation of JAK-STAT by a constitutively active form of Hop strongly increases both enteroendocrine cells and Stat92E activity.**

(A–B) Adult *Drosophila* posterior midgut from Esg-Gal4, 10XSTAT-GFP; Su(H)-GBE-Gal80, TubGal80$^{ts}$ driver expressing UAS-mCD8:RFP (CTL) or RNAis against core Atg genes involved at different stages of autophagy or UAS-Hop.H for 10 days, in intestinal stem cells. Related to Figure 6. (A) Quantification of Re(2)P dots per 10XSTAT-GFP$^+$ ISC and progenitors. (B) Representative maximal projections. 10XSTAT-GFP labels cells with active Stat92E (green). Prospero antibody marks enteroendocrine cells (magenta), and DAPI stains nuclei. Scale bar 50 μm. Quantification of (left graph) the percentage of total Pros$^+$ enteroendocrine cells, (right graph) mean per intestine of 10XSTAT-GFP RFI per cell normalized to the control, RFI (relative fluorescence intensity).

Data information: N = 4 independent experiments from four independent crosses. Quantifications represent the mean ± SEM. (A) Each dot represents Ref(2)P puncta per cell. (B) Each dot represents an intestine. The Kruskal-Wallis test was used, followed by Dunn's comparison tests. * $p<0.05$, *** $p < 0.001$, **** $p < 0.0001$, ns non-significant $p > 0.05$.

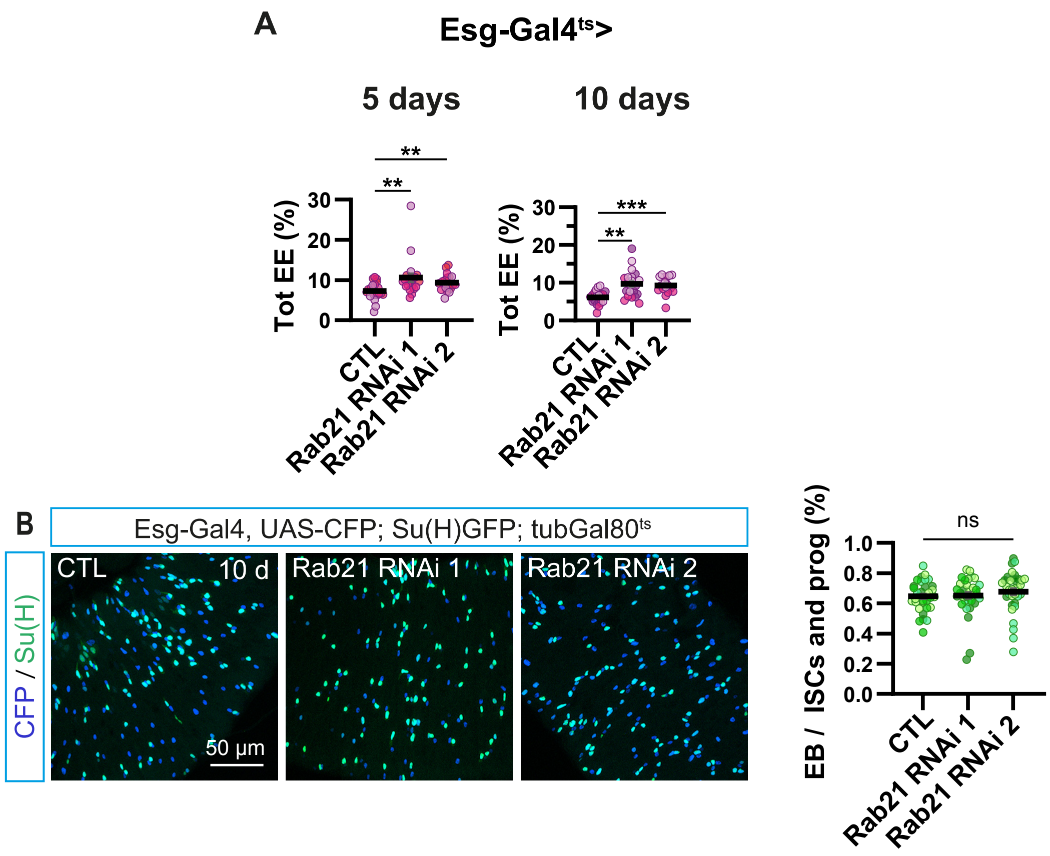


Supplementary Figure 1

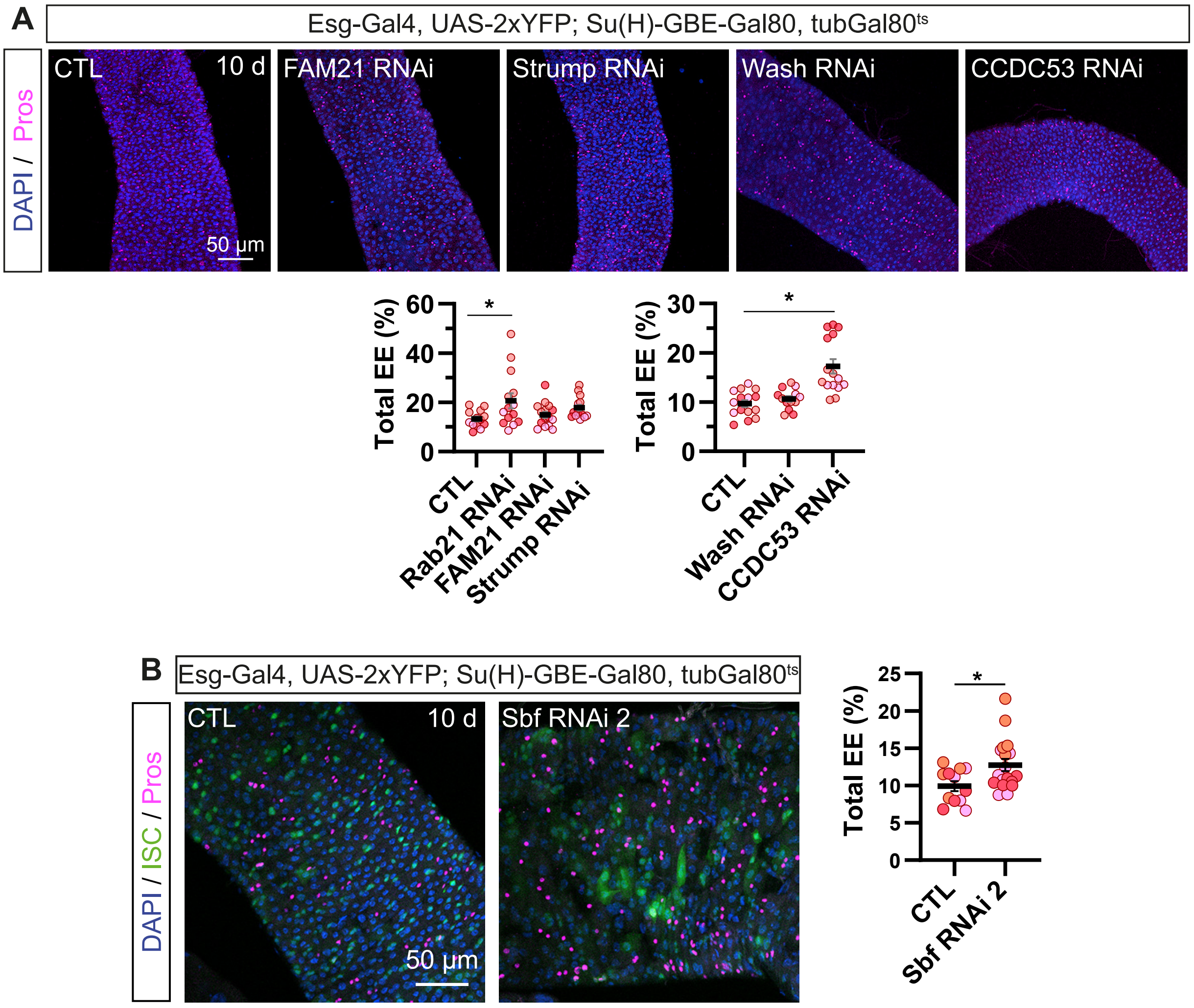


Supplementary Figure 2

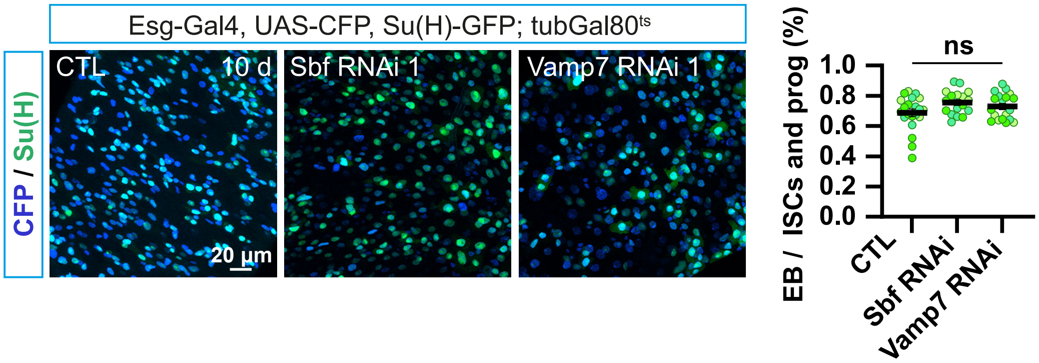


Supplementary Figure 3

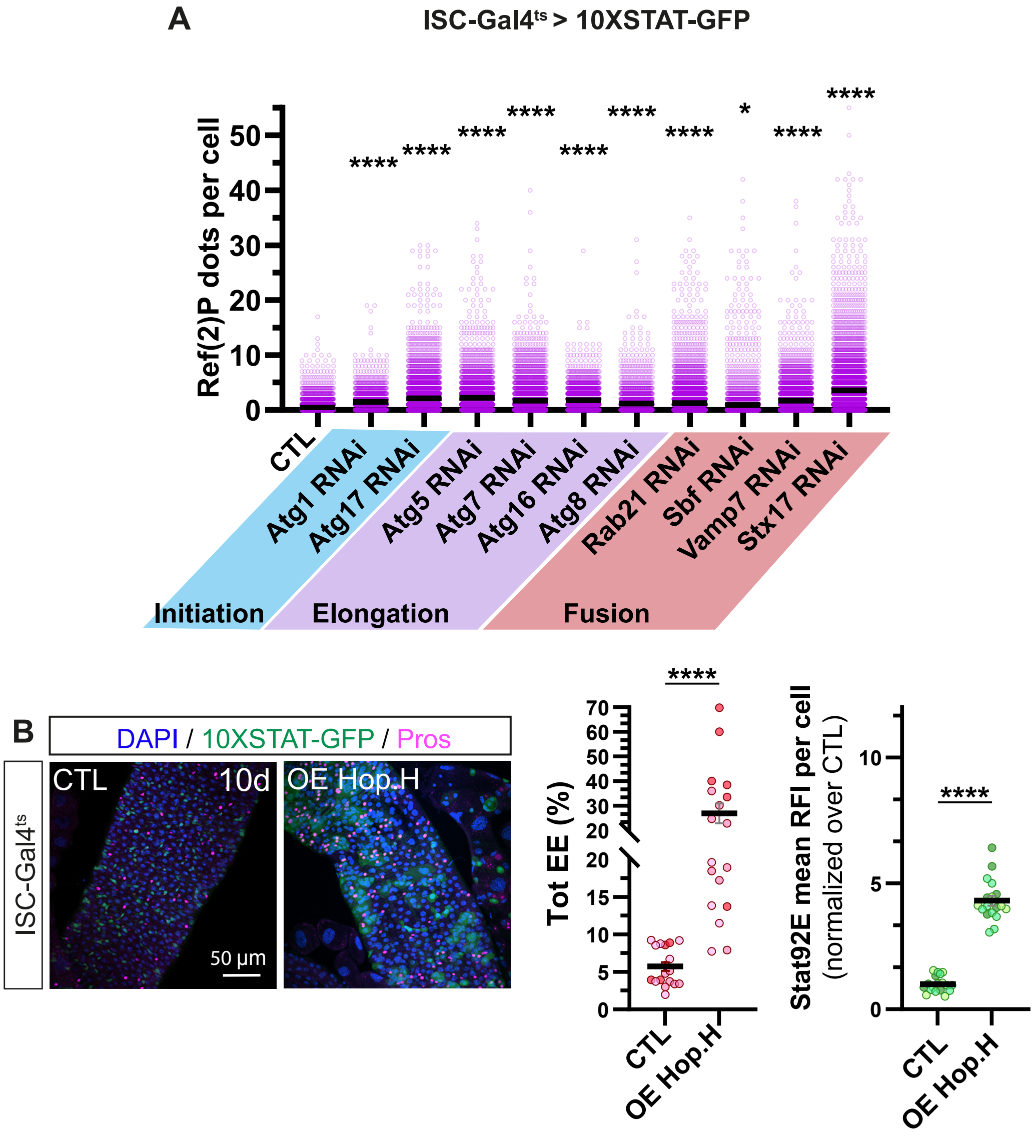


Supplementary Figure 4
